# Serum metabolomics signatures after an acute bout of combined traditional or high-intensity tactical training in young males and females

**DOI:** 10.1101/2024.08.16.608249

**Authors:** Zachary A. Graham, Khyatiben V. Pathak, Krystine Garcia-Mansfield, Kaleen M. Lavin, Anakaren R Torres, Jeremy S. McAdam, Timothy Broderick, Patrick Pirrotte, Marcas M. Bamman

**Affiliations:** Healthspan, Resilience and Performance, Florida Institute for Human and Machine Cognition, Pensacola, FL; Center for Exercise Medicine, The University of Alabama at Birmingham, Birmingham, AL; Department of Cell, Developmental and Integrative Biology, The University of Alabama at Birmingham, Birmingham, AL; Research Service, Birmingham VA Health Care System, Birmingham, AL; Early Detection and Prevention Division, Translational Genomics Institute, Phoenix, AZ; Integrated Mass Spectrometry Shared Resource, Translational Genomics Institute/City of Hope Comprehensive Cancer Center, Duarte, CA; Neurogenomics Division, Translational Genomics Institute, Phoenix, AZ

## Abstract

Exercise is a multipotent stimulus that results in large-scale dynamic changes to the systemic molecular profile. Alternative exercise prescriptions and doses would be expected to result in distinct signatures due to differences in duration and intensity. We tested two novel combined endurance and resistance exercise regimens to better understand how differing prescriptions alter the acute metabolomics response at multiple timepoints up to 24h post-exercise. Serum metabolomics for n=37 untrained individuals was analyzed for participants completing traditional combined exercise [TRAD; n = 20 (11M/9F)] or high-intensity tactical training [HITT; n= 17 (8M/9F)] before exercise (pre), and immediately (h0), 3 and 24 h post-exercise (h3 and h24, respectively). We found minimal metabolites had a group by time interaction (2 with FDR < 0.10; 31 with nominal p < 0.05;), but both stimuli resulted in large-scale within-group changes to the circulating metabolome. TRAD consistently had greater numbers of differentially abundant metabolites (FDR < 0.10) as compared to HITT at h0 (431 vs. 333), h3 (435 vs. 331) and h24 (168 vs. 76). The major metabolite classes altered were related to key energy substrates for both groups at h0 (e.g., glucose, pyruvate) and energy replenishment for h3 and h24 (e.g., 12,13 diHOME, palmitoylcarnitine, free fatty acids). In summary, our data are the first to describe the acute changes in the circulating metabolome following combined endurance and resistance exercise. Additionally, we show the two distinct doses of combined exercise led to generally similar patterns of responses, with the longer duration TRAD dose resulting in a higher magnitude of change.

## Introduction

Exercise improves health through a coordinated physiological response in almost all tissue systems and cell types. Long term adaptation to exercise training prevents or slows the onset of many diseases and improves health and function in those with existing pathologies (1). The diverse cues initiated by exercise are required to meet the demands of the activity while providing the molecular framework for protective long-term changes in cellular architecture that builds physiological reserve.

The metabolic demands of working tissue increases relative oxygen consumption by ∼25-fold during maximal exercise (2). Performing at this peak or at sustained submaximal levels requires elaborate inter- and intracellular communication. Increased accessibility to molecular profiling has greatly improved how we understand biological change at the molecular level (3). Metabolomics in particular is experiencing rapid advancements as sensitivity and analyte coverage has improved resolution, allowing for discovery of novel metabolite-specific mechanisms such as Lac-Phe (4). Thus, metabolomics allows further understanding of the molecular interplay required to maintain exercise performance, with unique signatures depending on mode and dose (5, 6).

Metabolomics has been performed across acute and chronic exercise studies (7), though most has been following acute endurance exercise (5). US public health guidelines include both endurance and resistance training as integral to a healthy lifestyle (8) and no timecourse study has investigated the acute circulating metabolomic response to combined endurance and resistance exercise. Recently, high-intensity interval exercise has been adopted as an alternative to traditional steady-state exercise, in particular in military personnel (9). This study aimed to address these gaps by investigating the circulating metabolome following an acute bout of traditional combined training (TRAD) or high-intensity tactical training (HITT).

## Methods

### Participants

Participants were recruited from the Birmingham, AL metro area as part of a larger parent trial (10), with this cohort previously described (11). Young, sedentary females and males completed a single acute bout of TRAD (n=23; 13M/9F) or HITT (n=17; 9M/8F). All individuals provided written, informed consent with protocol approval by the University of Alabama-Birmingham Institutional Review Board (#F160512012). Screening and enrollment procedures have been described (10).

### Exercise bout and blood collection

Exercise was completed in the early morning after an overnight fast. Upon arrival, participants rested supine for 30 min with blood collected via venipuncture. Exercise started ∼15 min post-blood draw. HITT performed 10 rounds of a maximal intensity 30 s ‘On/Off’ program of box jumps, burpees, split squat jumps, kettlebell swings, 2 cycling sprints, a rowing sprint, battle ropes, wall balls, and dips. TRAD performed 30 min of treadmill running at 70% heart rate reserve (HRR) calculated from VO_2peak_. Both groups then performed whole-body resistance training comprised of squats, knee extensions, heel raises, chest presses, overhead presses, seated rows, lat pulldowns, triceps pushdowns, biceps curl and abdominal crunches. TRAD completed 3×13 reps at 13 rep max with ∼60 s rest. HITT completed the same exercises in superset form at 3×9 reps at 9 rep max with 30 s rests between supersets. Exercise time was ∼90 min for TRAD and ∼45 min for HITT. Blood was collected immediately post-exercise (h0) then participants consumed a protein drink (350 kcal; 11 g fat/16 g protein/47 g carbohydrate). Blood was collected 3 h (h3) and 24 h post-exercise (fasted; h24). Blood was processed for serum by clotting at room temperature for 30 min then centrifuging at 2,000 g for 10 min at 4ºC. Serum was aliquoted and frozen at -80ºC. Sample sizes differ slightly compared to our previous report (11) because of serum availability. TRAD: n=20 at ‘pre’, n=17 at ‘h0’, n=16 at ‘h3’ and n=18 at ‘h24’; HITT: n=16 at ‘pre’, n=14 at ‘h0’, n=14 at ‘h3’ and n=16 at ‘h24’.

### Serum processing and mass spectrometry

Serum metabolites were extracted using a 1:3 plasma:organic solvent (3:1 v/v methanol:acetonitrile) ratio. Solvent was spiked with four isotopically labelled internal standards (d8-valine, ^13^C6-phenylalanine, ^13^C6-adipic acid, d4-succinic acid), vortexed for 30 s, then incubated at -20°C for 10 min for protein precipitation. Supernatant metabolites were recovered by centrifugation at 15,000g for 10 min at 4°C and subjected to untargeted metabolomics as described (12). 10 µl/sample was pooled for QA/QC and was injected prior to sample batches for system conditioning and every five samples to monitor instrument performance, extraction efficiency, correct retention time drift and perform batch normalization (13). Raw data were subjected to Compound Discoverer 3.2 for metabolite annotation and relative quantification (14).

### Data processing and statistical analysis

Data were processed and visualized with R (v3.6)(15) using UpSet (16), pheatmap (17) and mixOmics (18) packages. Metabolite abundances were analyzed using *limma* (19). Benjamini-Hochberg false discovery rate (FDR) of < 0.100 was the threshold. One participant failed outlier detection (6 > median absolute deviations; TRAD Male) and one participant was removed due to lack of ‘pre’ sample (HITT Female). The data matrix, metadata, and statistical outcomes are in Supplemental File 1.

## Results

### Participant Characteristics

General participant descriptions were previously published (11). The groups had no differences in age, height, weight, body composition, maximal aerobic (VO_2peak_) and anaerobic (Wingate) power, or strength.

### Metabolites

We detected 10,793 compounds across reverse phase and HILIC chromatographies in both positive and negative modes (Fig. 1A). 5,215 were annotated, with 3,243 non-redundant. 3,126 total compounds had FDR < 0.100 in at least one comparison, including 784 annotated endogenous metabolites belonging to major metabolite classes (Fig. 1B). PLS-DA plots show unique clustering for both groups at h0 and h3 (Fig. 1C) but not pre or h24.

**Fig. 1.**
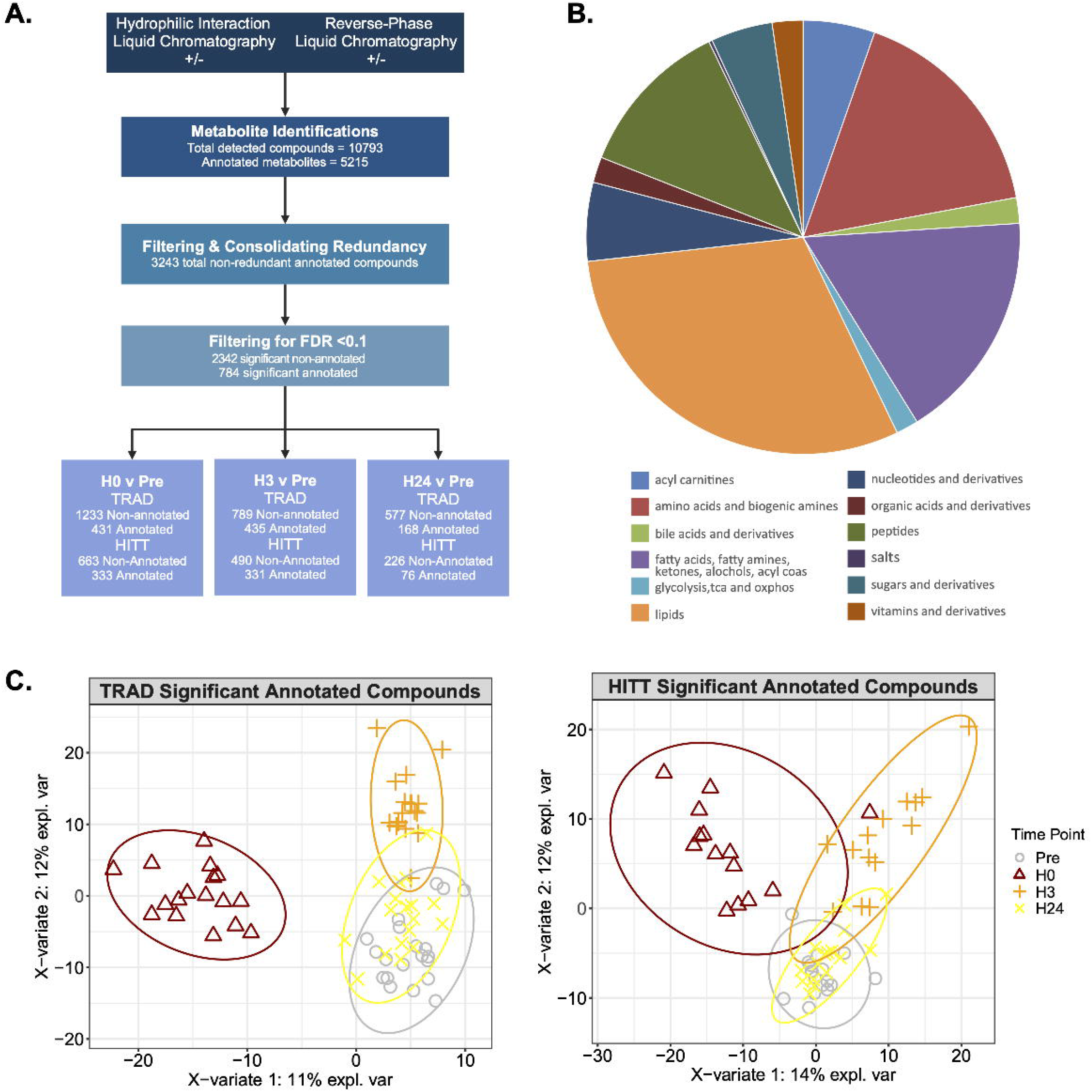
Serum metabolome identification. A) Summary of metabolomics profiling including total compounds, annotated and unannotated compounds, metabolite verification and consolidating redundant metabolites. B) Distribution of biochemical classes for 784 non-redundant significant metabolites across comparisons (h0vspre, h3vpre, h24vspre) between TRAD vs HITT. C) Partial least squares-discriminant analysis (PLS-DA) of differentially abundant metabolites (FDR < 0.10) across time for TRAD (lower left panel) and HITT (lower right panel), respectively.

The following analyses are on annotated metabolites only.

### Between groups comparisons

Comparisons of TRAD vs. HITT resulted in 31 metabolites with nominal p values < 0.010 (‘h0v pre’: 9; ‘h3vpre’: 18; ‘h24vpre’: 4) and are visualized via heatmap and UpSet plot (Fig. 2B). Only 2 had an FDR < 0.100: valine (FDR = 0.021) and capryloylglycine (FDR = 0.022), both lower in TRAD at ‘h0’ (Fig. 2C red squares). Fig. 2C shows boxplots for each metabolite with p < 0.010.

**Fig. 2.**
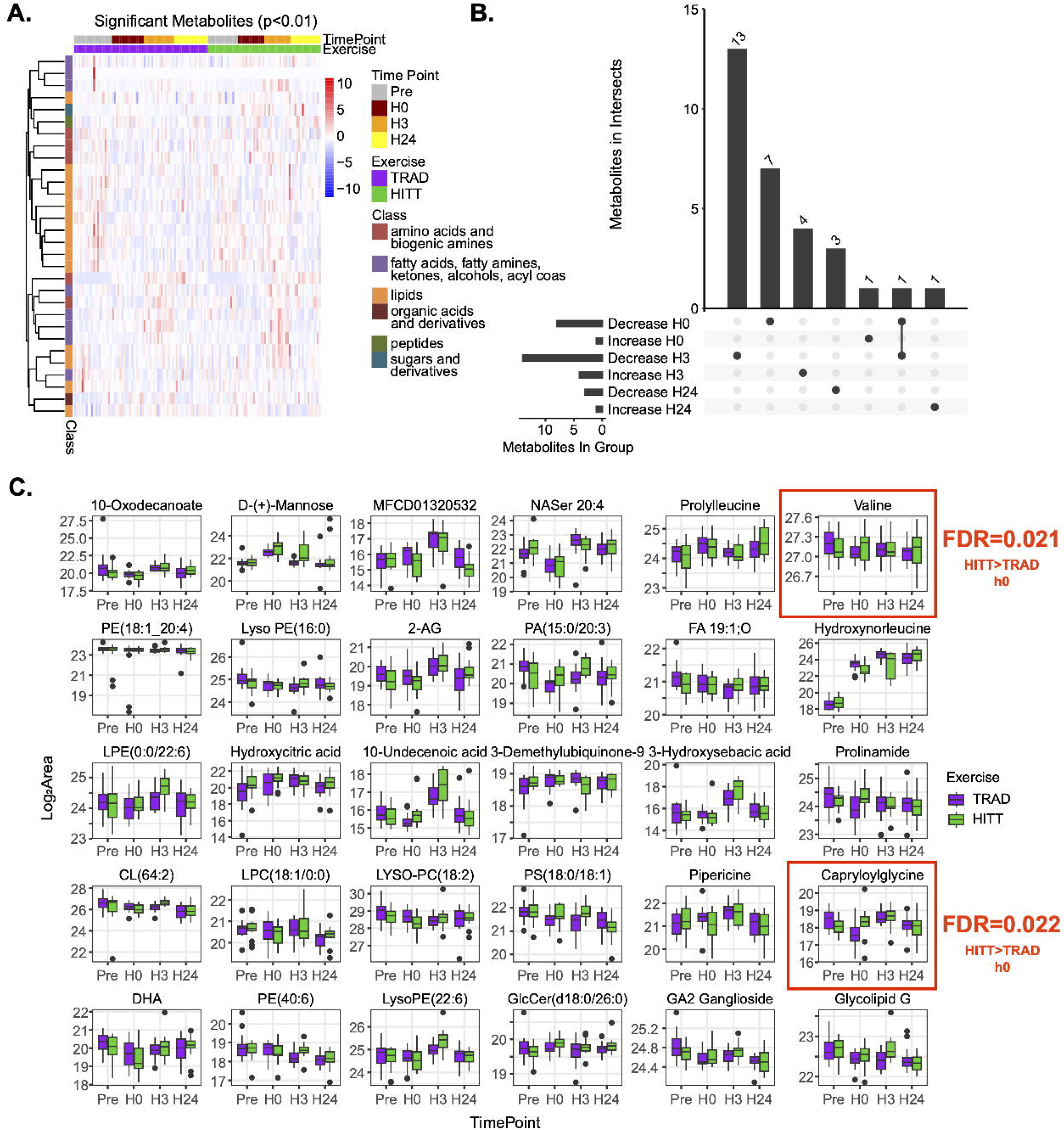
Differential circulating metabolomic responses for TRAD and HITT. A) Hierarchical clustering of differentially abundant metabolites by nominal p value (p < 0.01). B) Upset plot showing the patterns of time and direction response and C) box and whisker plots log showing log-transformed values for each metabolite per group and per timepoint. Metabolites highlighted with a red border are metabolites with FDR < 0.10.

### Within-group comparisons

Almost all 784 detected annotated metabolites showed a change postexercise: 667 and 552 metabolites were differentially abundant (FDR < 0.100) in TRAD and HITT, respectively (Fig. 3A). 431 metabolites (↑207, ↓224) were differentially abundant in TRAD at h0 compared to 333 (↑174, ↓159) for HITT. At h3, 435 metabolites (↑328, ↓107) and 331 metabolites (↑273, ↓58) were altered in TRAD and HITT, respectively. At h24, 168 (↑77, ↓91) metabolites were altered following TRAD and 76 (↑37, ↓39) following HITT.

**Fig. 3.**
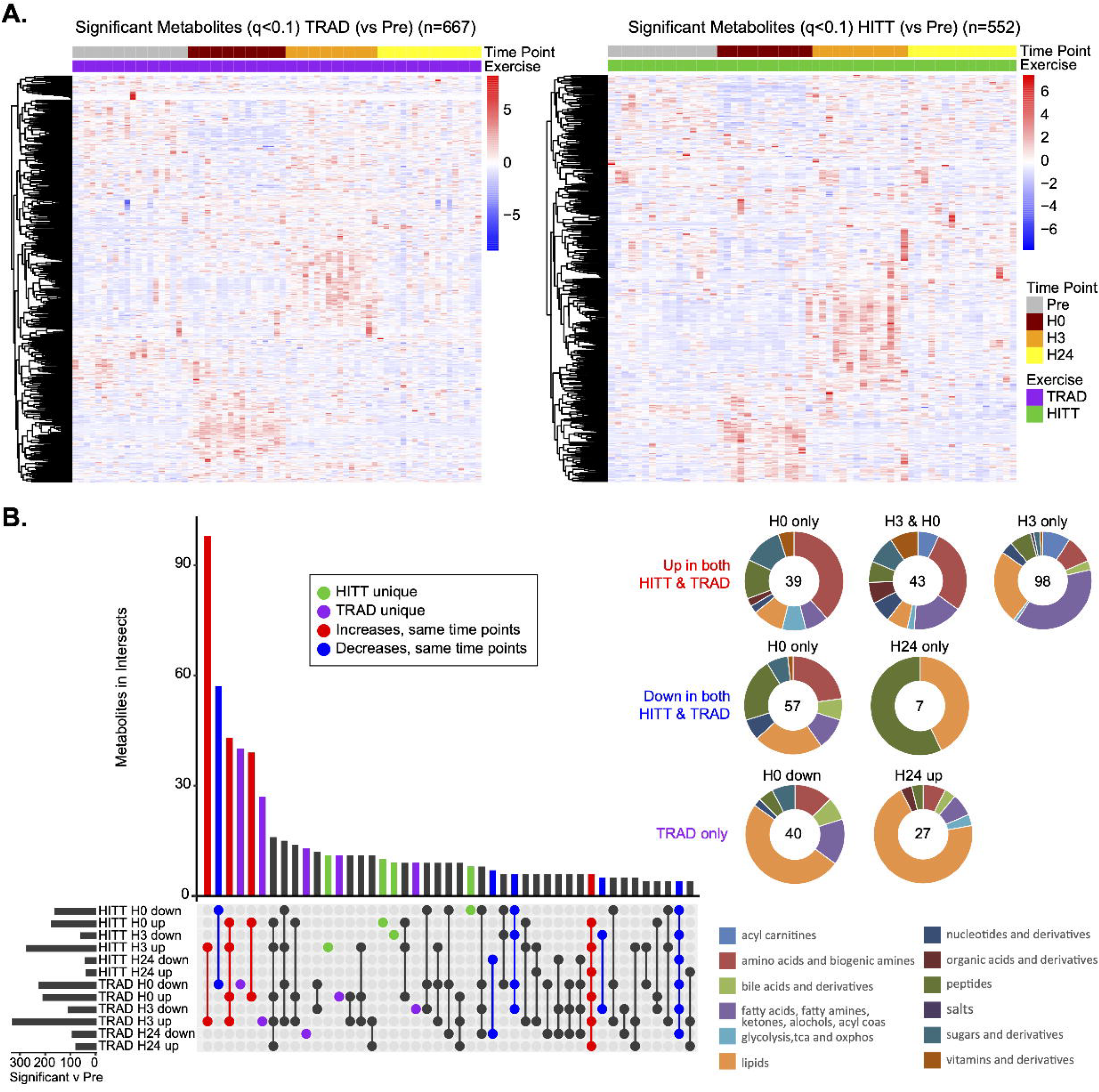
Within group comparisons across the exercise doses. A) Hierarchical clustering of differentially abundant metabolites (FDR < 0.10) for TRAD (Upper left panel) and HITT (Upper right panel). B) Upset plot representing shared or unique metabolite signatures for TRAD and HITT alongside donut charts representing biochemical classes.

Metabolite signatures and biological class membership are shown in Fig. 3B, with a focus on the 6 intersections with >25 metabolites. The ‘h3 up’ profile had the most shared metabolites with 98 metabolites mostly related to lipid and fatty acid metabolism common to both groups. Both groups shared 57 downregulated metabolites at ‘h0’ and annotated to amino acids and biogenic amines, lipids, and peptides. ‘h0 up’ and ‘h3 up’ had 43 metabolites shared and was composed of biogenic amines, acyl carnitines, fatty acids and derivatives. A group of 40 metabolites was only decreased in TRAD at ‘h0’ and comprised phospholipids, fatty acids and derivatives, and amino acids. At h0 both groups shared 39 upregulated metabolites, mostly amino acids and biogenic amines. Lastly, a set of 27 metabolites was increased only in TRAD at ‘h3’ largely annotated to phospholipids.

### Differential abundance patterns across time

Unbiased clustering was used to determine patterns of change relative to ‘pre’ (cluster membership ≥10 metabolites; Fig. 4A-B). Most signatures were shared between groups and are summarized in Table 1. There was a range of responses, from immediate increase (TRAD Cluster 1, HITT Cluster 3;) or decrease (TRAD Cluster 5, HITT Cluster 7) with a return to baseline a h24, immediate and sustained decrease (TRAD Cluster 6, HITT Cluster 6), or a unique signature such as HITT Cluster 8 which had stable resting and h0 abundances, but gradually increased at h3 and h24.

**Fig. 4.**
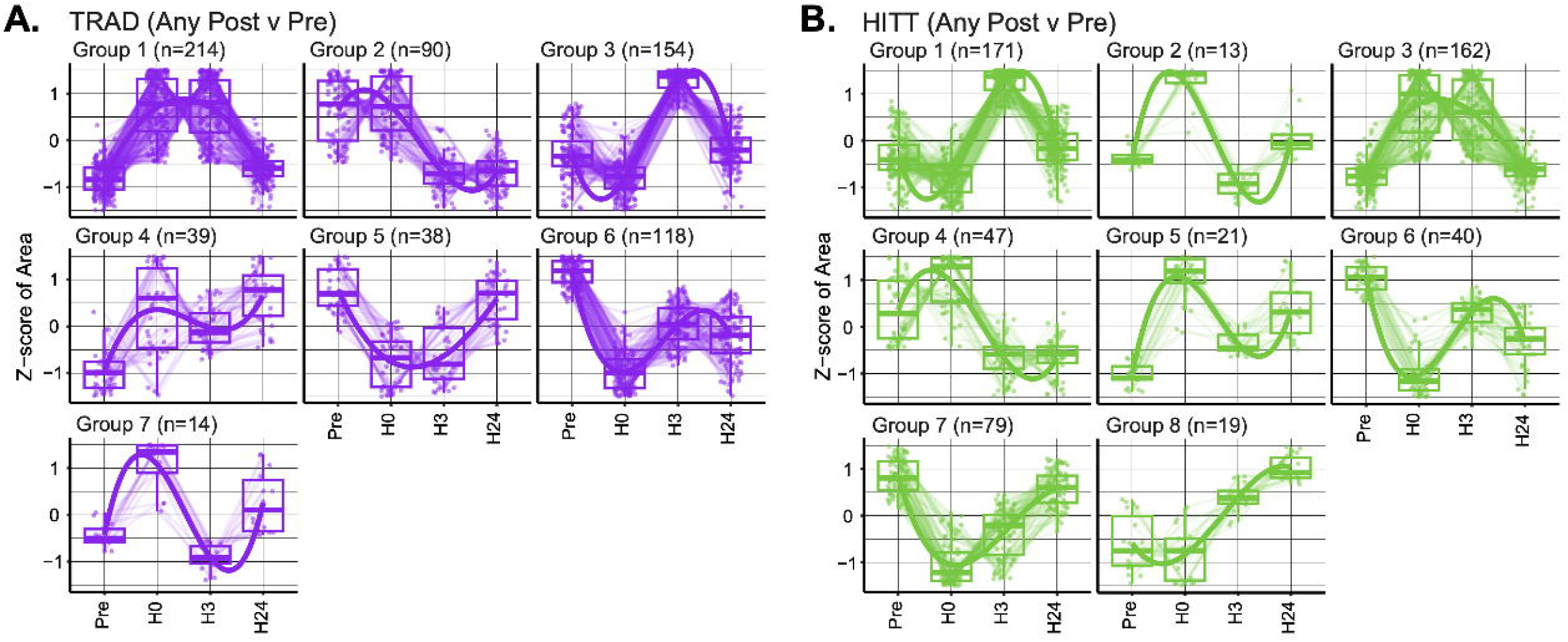
Unbiased clustering of all differential abundant metabolites compared to ‘pre’ for A) TRAD and B) HITT.

**Table 1.** Unbiased clustering of differentially abundant metabolites across time.

| Shared |  | Any timepoint vs. 'pre' |  |  |  |
| --- | --- | --- | --- | --- | --- |
| TRAD | HITT | Shared between Clusters | Shared % TRAD | Shared % HITT | Key metabolites |
| Cluster 1 | Cluster 3 | 117 | 55% | 72% | Glucose, aconitate, pyruvate |
| Cluster 2 | Cluster 4 | 27 | 30% | 57% | Cortisol, tryptophan, lactate |
| Cluster 3 | Cluster 1 | 97 | 63% | 58% | 12,13 di-HOME, palmitoylcarnitine, oleic acid |
| Cluster 4 | Cluster 5 | 7 | 18% | 33% | Glutathione |
| Cluster 5 | Cluster 7 | 18 | 47% | 23% | Succinate, thymidine, asparagine |
| Cluster 6 | Cluster 6 | 13 | 11% | 33% | Phenylalanine, 1-hydroxy-L-tryptophan |
| Cluster 7 | Cluster 2 | 0 | 0% | 0% | N/A |

## Discussion

Moderate, continuous endurance exercise has been the predominant focus of metabolomics-based acute exercise research (5). While others have demonstrated efficacy of subacute and chronic combined exercise training to change the resting metabolome (7), this is the first study to investigate how an acute bout of combined exercise alters the circulating metabolome and how different combinatorial exercise prescriptions impact this response. We identified 2 metabolites that met FDR criteria between TRAD and HITT, both downregulated in TRAD at h0: the branched-chain amino acid (BCAA) valine and the amino acid metabolite capryloylglycine. Valine is an important BCAA that can be used as an upstream substrate for the TCA intermediary succinyl-CoA (20). Serum valine levels are reduced following 80 min of sustained treadmill walking in males exposed to lab-based high-altitude equivalency (21) and following 80 min of sustained moderate cycling (∼65% VO_2_max) in males with adequate carbohydrate intake; though those with calorically-matched, low-carbohydrate intake saw elevated abundances of valine (22). TRAD was roughly 90 min and more sustained than HITT, and our data mostly supports long-duration exercise reduces circulating valine. Capryloylglycine is a medium-chain fatty acid conjugate often used in cosmetic products. However, it has been described as a metabolite in mammalian systems. In pigs, capryloylglycine is a principal metabolite produced by the pancreas (23) and in humans it is elevated in skeletal muscle biopsies of older adults compared to younger adults (24). Genetically-induced mitochondrial β-oxidation deficiencies lead to elevated capryloylglycine levels that are excreted in urine (25). Why capryloylglycine was reduced immediately after TRAD is unknown as glycine levels were similar across both groups, suggesting equal availability. It is possible levels of caprylic acid were lower but it was not among our annotated metabolites. Another possibility is TRAD reduces activity of glycine N-acyltransferase, the enzyme that conjugates glycine to fatty acids. Future investigations are needed to resolve whether this change is biologically impactful.

TRAD had consistently more metabolites with differential abundance at each timepoint compared to HITT in our within-group analyses. This is in line with our transcriptomics work from this same cohort that reported generally elevated numbers of differentially expressed RNA species in TRAD vs. HITT (e.g., long non-coding RNA, protein coding RNA, microRNA) in serum-derived extracellular vesicles and skeletal muscle (11). This is likely due to a combination of factors such as the sustained metabolic demand from the continuous nature of the endurance portion as well as the greater length of exercise duration for the TRAD group (∼90 min) compared to HITT (∼45 min). Despite the ‘h0’ timepoint being a unique cluster as identified by PLS-DA in both TRAD and HITT as shown in our PLS-DA plot, the ‘h0’ and ‘h3’ timepoints had roughly equivalent numbers of metabolites for each group. There was a differential pattern as ‘h0’ had roughly identical numbers of upregulated and downregulated metabolites while ‘h3’ was composed largely of upregulated metabolites. The upregulated metabolites in ‘h3’ were mostly annotated to fatty acids, conjugated fatty amines, and ketones. These represented classes fall in-line with systemic replenishment of energy stores that is supported by both TRAD and HITT having elevated levels of the adipose tissue-released exerkine 12,13 di-HOME, which increases fatty acid uptake in skeletal muscle (26), elevated levels of palmitoylcarnitine, which is derived from palmitoyl-CoA and shuttled across the inner membrane of the mitochondria (where it is subsequently converted back into palmitoyl-CoA for β-oxidation), and a host of other long chain fatty acids and their metabolites. The ‘h0’ timepoint was more associated with the active substrates of ATP generation (glucose, pyruvate, citrate, aconitate, fumarate) and the purine catabolism/salvage pathways (purine, guanine, hypoxanthine, uridine monophosphate), and is supported by unimodal studies using 45-60 min of moderate to moderate-vigorous steady state cycling or traditional resistance training at similar post-exercise timepoints (6, 27). Purine metabolism and ultimate salvage is more energy-efficient than de novo formation to maintain ATP levels, but the elevation in the blood levels of these metabolites would be expected to be from working skeletal muscle. Xanthine oxidase, the enzyme that converts hypoxanthine to uric acid, is not expressed in skeletal muscle so these metabolites are likely taken up by the liver to be salvaged or further metabolized to uric acid (28).

There were unique differences in metabolite signatures at ‘h24’ vs. ‘pre’ with TRAD having a more sustained profile. Of the 168 differentially abundant metabolites in TRAD at h24, 111 of them (68%) were also differentially abundant in our ‘h3vpre’ comparison. This was not the case with HITT, as only 43% (36/76) remained differentially abundant. Despite the differences in overall numbers and the proportion common at ‘h3’ and ‘h24’ between TRAD and HITT, amino acid metabolites, in particular the BCAAs leucine and isoleucine, were similarly elevated. This is contrasted by unimodal endurance or resistance training acute bouts that reduced BCAAs, though their metabolites were elevated in the immediate and short timecourse after exercise (0-120 min)(6, 22), which we did observe. Similar to TRAD having a more robust response in the number of differentially abundant metabolites, this continual change in metabolite levels is likely related to the longer, continuous duration of the TRAD dose.

Our combinatorial approach complicates direct comparisons against the literature as both TRAD and HITT were novel prescriptions. However, the magnitude of response was similar to both acute resistance and endurance bouts of exercise that saw peak changes immediately, 15 min, and out to 3 h post-exercise (6). In both groups, the resistance portion was completed last. Thus, there is potential that our metabolomic profiling would more mimic a resistance training prescription, however, the relative TRAD and HITT up and downregulated proportions at h0 and h3 timepoints do not align with resistance and endurance unimodal proportions at these same timepoints (6), suggesting our combinatorial intervention results in a unique metabolomic dynamic.

Our study has a few considerations. For sex as a biological variable, our randomization strategy included stratification by sex to ensure relative balance between HITT and TRAD, but we did not covary or normalize for sex similar to our approach with the transcriptome (11). Importantly, however, we have recently reported the sex-specific multiomic responses for these participants, including the metabolome (29). Another important consideration is the supplemental protein drink after the ‘h0’ blood draw. This was used to mimic a real-world exercise scenario and we cannot rule out that some of the shared metabolites observed to be changed at ‘h3’, or less likely, ‘h24’ between both groups were from the supplement. However, this concern is dampened by evidence showing our general patterns of change in classes such as lipids and amino acids match those from both endurance and resistance training acute response studies where participants remained fasted for serial blood analyses over 3h post-exercise (6). In summary, our novel combined exercise prescriptions resulted in robust metabolomic changes across multiple timepoints post-acute exercise that share major characteristics of unimodal exercise bouts, but also suggest distinct signatures related to exercise dose.

## Acknowledgements

This study was funded by Department of Defense Office of Naval Research Grant N000141613159 (MMB, TJB). Additional support was provided by the Department of Veterans Affairs Rehabilitation Research and Development Service Grant IK2RX002781 (ZAG) and the National Institutes of Health Grant P30CA033572 (PP) The views represented in this manuscript are not reflective of the United States Government or the Department of Veterans Affairs.

We thank the participants for volunteering and completing the exercise bouts and the research and exercise training staff for their time and effort in helping successfully execute the study.

## References

1. Pedersen BK, Saltin B. Exercise as medicine - evidence for prescribing exercise as therapy in 26 different chronic diseases. Scand J Med Sci Sports 25 Suppl 3: 1–72, 2015. doi: 10.1111/sms.12581.

2. Jones AM, Kirby BS, Clark IE, Rice HM, Fulkerson E, Wylie LJ, Wilkerson DP, Vanhatalo A, Wilkins BW. Physiological demands of running at 2-hour marathon race pace. J Appl Physiol Bethesda Md 1985 130: 369–379, 2021. doi: 10.1152/japplphysiol.00647.2020.

3. Sanford JA, Nogiec CD, Lindholm ME, Adkins JN, Amar D, Dasari S, Drugan JK, Fernández FM, Radom-Aizik S, Schenk S, Snyder MP, Tracy RP, Vanderboom P, Trappe S, Walsh MJ, Molecular Transducers of Physical Activity Consortium. Molecular Transducers of Physical Activity Consortium (MoTrPAC): Mapping the Dynamic Responses to Exercise. Cell 181: 1464–1474, 2020. doi: 10.1016/j.cell.2020.06.004.

4. Li VL, He Y, Contrepois K, Liu H, Kim JT, Wiggenhorn AL, Tanzo JT, Tung AS-H, Lyu X, Zushin P-JH, Jansen RS, Michael B, Loh KY, Yang AC, Carl CS, Voldstedlund CT, Wei W, Terrell SM, Moeller BC, Arthur RM, Wallis GA, Van De Wetering K, Stahl A, Kiens B, Richter EA, Banik SM, Snyder MP, Xu Y, Long JZ. An exercise-inducible metabolite that suppresses feeding and obesity. Nature 606: 785–790, 2022. doi: 10.1038/s41586-022-04828-5.

5. Schranner D, Kastenmüller G, Schönfelder M, Römisch-Margl W, Wackerhage H. Metabolite Concentration Changes in Humans After a Bout of Exercise: a Systematic Review of Exercise Metabolomics Studies. Sports Med - Open 6: 11, 2020. doi: 10.1186/s40798-020-0238-4.

6. Morville T, Sahl RE, Moritz T, Helge JW, Clemmensen C. Plasma Metabolome Profiling of Resistance Exercise and Endurance Exercise in Humans. Cell Rep 33: 108554, 2020. doi: 10.1016/j.celrep.2020.108554.

7. Jaguri A, Al Thani AA, Elrayess MA. Exercise Metabolome: Insights for Health and Performance. Metabolites 13: 694, 2023. doi: 10.3390/metabo13060694.

8. Piercy KL, Troiano RP, Ballard RM, Carlson SA, Fulton JE, Galuska DA, George SM, Olson RD. The Physical Activity Guidelines for Americans. JAMA 320: 2020–2028, 2018. doi: 10.1001/jama.2018.14854.

9. Haddock CK, Poston WSC, Heinrich KM, Jahnke SA, Jitnarin N. The Benefits of High-Intensity Functional Training Fitness Programs for Military Personnel. Mil Med 181: e1508–e1514, 2016. doi: 10.7205/MILMED-D-15-00503.

10. McAdam JS, Craig MP, Graham ZA, Peoples B, Tuggle SC, Seay RS, Lavin KM, Sullivan AB, O’Bryan SM, Yang S, Drummer DJ, Kelley CJ, Peri K, Bell MB, Aban I, Cutter GR, Mahyari A, Wen Y, Zhang J, Hira A, Broderick TJ, Kadakia M, Bamman MM. Biocircuitry linked to exercise adaptations: impact of dose and inter-individual response heterogeneity. Sports Medicine: 2025.

11. Lavin KM, Graham ZA, McAdam JS, O’Bryan SM, Drummer D, Bell MB, Kelley CJ, Lixandrão ME, Peoples B, Tuggle SC, Seay RS, Van Keuren-Jensen K, Huentelman MJ, Pirrotte P, Reiman R, Alsop E, Hutchins E, Antone J, Bonfitto A, Meechoovet B, Palade J, Talboom JS, Sullivan A, Aban I, Peri K, Broderick TJ, Bamman MM. Dynamic transcriptomic responses to divergent acute exercise stimuli in young adults. Physiol Genomics 55: 194–212, 2023. doi: 10.1152/physiolgenomics.00144.2022.

12. Zhang B, Zhao D, Chen F, Frankhouser D, Wang H, Pathak KV, Dong L, Torres A, Garcia-Mansfield K, Zhang Y, Hoang DH, Chen M-H, Tao S, Cho H, Liang Y, Perrotti D, Branciamore S, Rockne R, Wu X, Ghoda L, Li L, Jin J, Chen J, Yu J, Caligiuri MA, Kuo Y-H, Boldin M, Su R, Swiderski P, Kortylewski M, Pirrotte P, Nguyen LXT, Marcucci G. Acquired miR-142 deficit in leukemic stem cells suffices to drive chronic myeloid leukemia into blast crisis. Nat Commun 14: 5325, 2023. doi: 10.1038/s41467-023-41167-z.

13. Dunn WB, Broadhurst D, Begley P, Zelena E, Francis-McIntyre S, Anderson N, Brown M, Knowles JD, Halsall A, Haselden JN, Nicholls AW, Wilson ID, Kell DB, Goodacre R, Human Serum Metabolome (HUSERMET) Consortium. Procedures for large-scale metabolic profiling of serum and plasma using gas chromatography and liquid chromatography coupled to mass spectrometry. Nat Protoc 6: 1060–1083, 2011. doi: 10.1038/nprot.2011.335.

14. Najdekr L, Blanco GR, Dunn WB. Collection of Untargeted Metabolomic Data for Mammalian Urine Applying HILIC and Reversed Phase Ultra Performance Liquid Chromatography Methods Coupled to a Q Exactive Mass Spectrometer. Methods Mol Biol Clifton NJ 1996: 1–15, 2019. doi: 10.1007/978-1-4939-9488-5_1.

15. R-Core-Team. R: A Language and Environment for Statistical Computing [Online]. R Foundation for Statistical Computing: 2014. http://www.R-project.org.

16. Conway JR, Lex A, Gehlenborg N. UpSetR: an R package for the visualization of intersecting sets and their properties. Bioinforma Oxf Engl 33: 2938–2940, 2017. doi: 10.1093/bioinformatics/btx364.

17. Kolde R. Pheatmap: pretty heatmaps. R Package Version 1: 726, 2019.

18. Rohart F, Gautier B, Singh A, Le Cao KA. mixOmics: An R package for ‘omics feature selection and multiple data integration. PLoS Comput Biol 13: e1005752, 2017. doi: 10.1371/journal.pcbi.1005752.

19. Law CW, Zeglinski K, Dong X, Alhamdoosh M, Smyth GK, Ritchie ME. A guide to creating design matrices for gene expression experiments. F1000Research 9: 1444, 2020. doi: 10.12688/f1000research.27893.1.

20. Adeva-Andany MM, López-Maside L, Donapetry-García C, Fernández-Fernández C, Sixto-Leal C. Enzymes involved in branched-chain amino acid metabolism in humans. Amino Acids 49: 1005–1028, 2017. doi: 10.1007/s00726-017-2412-7.

21. Margolis LM, Karl JP, Wilson MA, Coleman JL, Ferrando AA, Young AJ, Pasiakos SM. Metabolomic profiles are reflective of hypoxia-induced insulin resistance during exercise in healthy young adult males. Am J Physiol Regul Integr Comp Physiol 321: R1–R11, 2021. doi: 10.1152/ajpregu.00076.2021.

22. Margolis LM, Karl JP, Wilson MA, Coleman JL, Whitney CC, Pasiakos SM. Serum Branched-Chain Amino Acid Metabolites Increase in Males When Aerobic Exercise Is Initiated with Low Muscle Glycogen. Metabolites 11: 828, 2021. doi: 10.3390/metabo11120828.

23. Jang C, Hui S, Zeng X, Cowan AJ, Wang L, Chen L, Morscher RJ, Reyes J, Frezza C, Hwang HY, Imai A, Saito Y, Okamoto K, Vaspoli C, Kasprenski L, Zsido GA, Gorman JH, Gorman RC, Rabinowitz JD. Metabolite Exchange between Mammalian Organs Quantified in Pigs. Cell Metab 30: 594–606.e3, 2019. doi: 10.1016/j.cmet.2019.06.002.

24. Wilkinson DJ, Rodriguez-Blanco G, Dunn WB, Phillips BE, Williams JP, Greenhaff PL, Smith K, Gallagher IJ, Atherton PJ. Untargeted metabolomics for uncovering biological markers of human skeletal muscle ageing. Aging 12: 12517–12533, 2020. doi: 10.18632/aging.103513.

25. Costa CG, Guérand WS, Struys EA, Holwerda U, ten Brink HJ, Tavares de Almeida I, Duran M, Jakobs C. Quantitative analysis of urinary acylglycines for the diagnosis of beta-oxidation defects using GC-NCI-MS. J Pharm Biomed Anal 21: 1215–1224, 2000. doi: 10.1016/s0731-7085(99)00235-6.

26. Stanford KI, Lynes MD, Takahashi H, Baer LA, Arts PJ, May FJ, Lehnig AC, Middelbeek RJW, Richard JJ, So K, Chen EY, Gao F, Narain NR, Distefano G, Shettigar VK, Hirshman MF, Ziolo MT, Kiebish MA, Tseng Y-H, Coen PM, Goodyear LJ. 12,13-diHOME: An Exercise-Induced Lipokine that Increases Skeletal Muscle Fatty Acid Uptake. Cell Metab 27: 1111–1120.e3, 2018. doi: 10.1016/j.cmet.2018.03.020.

27. Thompson PD, Franklin BA, Balady GJ, Blair SN, Corrado D, Estes NA 3rd, Fulton JE, Gordon NF, Haskell WL, Link MS, Maron BJ, Mittleman MA, Pelliccia A, Wenger NK, Willich SN, Costa F. Exercise and acute cardiovascular events placing the risks into perspective: a scientific statement from the American Heart Association Council on Nutrition, Physical Activity, and Metabolism and the Council on Clinical Cardiology. Circulation 115: 2358–68, 2007. doi: 10.1161/circulationaha.107.181485.

28. Miller SG, Hafen PS, Brault JJ. Increased Adenine Nucleotide Degradation in Skeletal Muscle Atrophy. Int J Mol Sci 21: 88, 2019. doi: 10.3390/ijms21010088.

29. Lavin KM, O’Bryan SM, Pathak KV, Garcia-Mansfield K, Graham ZA, McAdam JS, Drummer DJ, Bell MB, Kelley CJ, Lixandrão ME, Peoples B, Seay RS, Torres AR, Reiman R, Alsop E, Hutchins E, Bonfitto A, Antone J, Palade J, Van Keuren-Jensen K, Huentelman MJ, Pirrotte P, Broderick T, Bamman MM. Divergent Multiomic Acute Exercise Responses Reveal the Impact of Sex as a Biological Variable..

